# Object speed and distractor number do not affect attentional allocation in multiple object tracking

**DOI:** 10.64898/2025.12.28.696734

**Authors:** N. Adamian, F. Akalan, S.K. Andersen

## Abstract

Keeping track of multiple moving objects across dynamic real-world scenarios such as driving, team sports, or crowded social environments is a fundamental challenge for visual attention. We have previously demonstrated that as the number of tracked objects increases, the strength of attentional facilitation allocated to each individual object decreases, limiting tracking success. It is also well established that beyond the number of tracked objects, faster-moving objects and objects embedded amongst higher numbers of distractors are more difficult to track. Are these effects on tracking difficulty also mediated by less effective allocation of attention to tracked targets as in the case of tracking more targets? If so, one should expect the strength of attentional modulation to drop systematically with increasing speed and total number of moving stimuli.

In two experiments (total n = 70), participants were instructed to track moving targets amongst identical distractors while we manipulated object speed (Experiment 1) and number (Experiment 2). As expected, tracking performance declined with both manipulations. However, steady-state visual evoked potentials (SSVEPs) recorded during successful tracking revealed that attentional enhancement of tracked targets compared with distractors did not drop with increasing speed or object number. In summary, bottom-up changes in the stimulus display and top-down attentional manipulations affect tracking performance in independent ways, with the balance between strength of attentional allocation and bottom-up demands of the task determining successful tracking. The allocation of attention itself seems to be determined exclusively by top-down goals rather than being reactive to bottom-up display characteristics.

**Open Practices Statement:** Participant level data and analysis code for all experiments are available at (https://osf.io/ypgfs/) and Experiment 1 was preregistered (https://osf.io/pxh25/).

**Significance statement:** Keeping track of multiple moving objects is fundamental to navigating dynamic real-world scenarios. This ability is accomplished through multifocal attentional selection, which weakens as the number of tracked targets increases. This study asks whether other stimulus manipulations increase tracking difficulty by diluting attentional allocation. Using steady-state visual evoked potentials to measure selective attention during tracking, we demonstrate that both increases in speed and distractor number impair performance, however, they do not affect attentional enhancement of targets. This suggests that top-down attentional control operates independently from bottom-up demands.

## Introduction

Human observers demonstrate a remarkable ability to visually track multiple independently moving objects, even when these targets are embedded amongst identical distractors (Holcombe, 2023; Pylyshyn & Storm, 1988). The mechanisms enabling us to simultaneously track multiple moving objects, and the fundamental limitations on this capacity, have been a central focus of research for decades (Meyerhoff et al., 2017).

The limits on tracking manifest across different aspects of the dynamic display. One of the most fundamental constraints is the limit on the number of objects that can be continuously tracked. While early research suggested that capacity was fixed at four objects (Pylyshyn & Storm, 1988), this theory was at odds with the discovery of a smooth trade-off between the number of tracked objects and their speed. When objects move slowly, participants can track up to as many as 8-9 objects, whereas at high speeds they may only be able to track a single one (Alvarez & Franconeri, 2007; Holcombe & Chen, 2012; Liu et al., 2005; Tombu & Seiffert, 2011). This finding was interpreted as evidence of a limited resource that is flexibly divided among the targets in line with task demands. The more difficult tracking gets, for example due to increased proximity of objects in the display, the more “resource” each target needs in order to be successfully tracked, which in turn limits tracking capacity (Iordanescu et al., 2009).

Numerous studies confirmed that tracking performance declines as object speed increases (Bettencourt & Somers, 2009; Drew et al., 2013; Feria, 2013; Tombu & Seiffert, 2011), although the underlying mechanisms remain debated. One possibility, in line with the aforementioned flexible resource model (Alvarez & Franconeri, 2007), is that faster moving objects increase attentional demands on tracking, and thus speed itself affects tracking performance (Feria, 2013; Holcombe & Chen, 2012; Tombu & Seiffert, 2011). Alternatively, it has been proposed that increased speed reduces tracking performance solely because it increases the number of close encounters between objects, leading to spatial interference and target-distractor confusions (Franconeri et al., 2008, 2010).

Another confirmed prediction of the spatial interference account of multiple object tracking is that for a fixed number of targets, tracking becomes more difficult when additional distractors are present (Bettencourt & Somers, 2009; Drew et al., 2013; Feria, 2012). This increases the likelihood of targets and distractors coming in close proximity and causing interference, which can be overcome through increasing attentional demands on tracking (Franconeri et al., 2010). However, Bettencourt and Somers (2009) proposed a different explanation – if distractors require active suppression, they take away from the attention available for targets, and thus tracking becomes more difficult with more distractors.

Overall, while there is consensus that tracking is flexible, capacity-limited, and relies on attention, the nature of this attentional resource and the ways in which flexibility manifests remain debated. A recent study from our lab (Adamian & Andersen, 2022) demonstrated that the magnitude of attentional amplification of tracked targets measured with steady-state visual evoked potentials (SSVEPs) decreased with increasing set size. Moreover, this strength of attentional selection was predictive of tracking success on a given trial. This suggests that the allocation of attention in early visual cortex limits how many objects can be simultaneously tracked. The present study extends this work by examining whether attentional amplification of targets depends on other factors known to affect tracking: object speed and the total number of objects present.

In two experiments, participants tracked moving objects among identical distractors. Targets and distractors flickered at different frequencies, each eliciting steady-state visual evoked potentials (SSVEPs) at their respective flicker frequencies. This frequency-tagging approach resulted in separable neural activity for each stimulus type, allowing us to concurrently measure the attentional facilitation of both targets and distractors throughout the tracking period. In Experiment 1, the speed of motion was manipulated while the number of targets and distractors was constant. In Experiment 2, the speed was constant as was the number of targets (four), however, the number of distractors was manipulated. If object speed or total number of objects affect how limited attention is distributed – for example, if faster speeds result in attention not ‘keeping up’ with motion or if extra items challenge the spatial resolution of attention – we expect to observe a decline in attentional selectivity corresponding to decline in tracking performance resulting from these manipulations.

## Experiment 1: Speed

### Method

Data collection and analysis of Experiment 1 were preregistered: https://osf.io/pxh25

#### Participants

A target sample size of 17 participants was determined through power analysis (detailed in the preregistration protocol). Twenty-two members of the University of Aberdeen student community participated in the study (6 men, all right-handed; M_age_ = 26 years, SD_age_ = 6.08). All participants provided written informed consent and received £20 for their time. Participants reported normal color vision and normal or corrected-to-normal visual acuity. The study received ethical approval from the Psychology Ethics Committee at the University of Aberdeen (PEC/5048/2022/7).

Data from five participants were excluded because fewer than 20 epochs remained in at least one experimental condition after rejecting trials with EEG artefacts and trials with incorrect behavioral responses, resulting in a final sample of 17 participants.

#### Stimuli and procedure

Stimuli were created using MATLAB (MathWorks Inc., Natick, MA) and Psychtoolbox (Kleiner et al., 2007). They were presented in a dimly lit room on a 32” Display++ monitor (CRS, UK) with a resolution of 1920×1080 and refresh rate of 120 Hz. Participants were seated at a viewing distance of approximately 60 cm without head restraint and instructed to maintain fixation on a central fixation point (0.3 dva). Stimuli were presented against a mid-grey background (29 cd/m²) within a centrally positioned light-grey elliptical field (38.5 cd/m², 36 dva width, 28 dva height). The display consisted of eight identical red discs (12.2 cd/m², 4 dva diameter). Target cues were provided by black outlines around each disc, whilst feedback was given by outlining cued discs in green (correct) or red (incorrect).

Each trial began with the eight red discs randomly positioned within the elliptical viewing area (see Figure 1). Four discs were then outlined as to-be-tracked targets. After 1250 ms the outlines disappeared and all objects began moving linearly in randomly chosen directions, bouncing off each other and the borders of the background field at physically realistic angles. Throughout both cueing and tracking, all objects flickered at their designated frequencies. To prevent discs from overlapping they were surrounded by an invisible “bumper” border 1 dva wider than the disc itself. Movement speed varied by condition: slow (3.8 dva/s), medium (7.6 dva/s), or high (10.4 dva/s). These speeds were selected to yield performance levels comparable to those observed when the number of tracked targets was manipulated in a previous study (Adamian & Andersen, 2022). After a 4000 ms tracking period, objects stopped moving and participants selected the discs they thought were targets via mouse clicks. Participants always selected exactly four discs, guessing when necessary. Visual feedback (green outlines for correct selections, red for incorrect) and auditory feedback (high-pitch tone for all correct, low-pitch otherwise) were provided after each trial. Summary feedback was also provided after each block.

**Figure 1.**
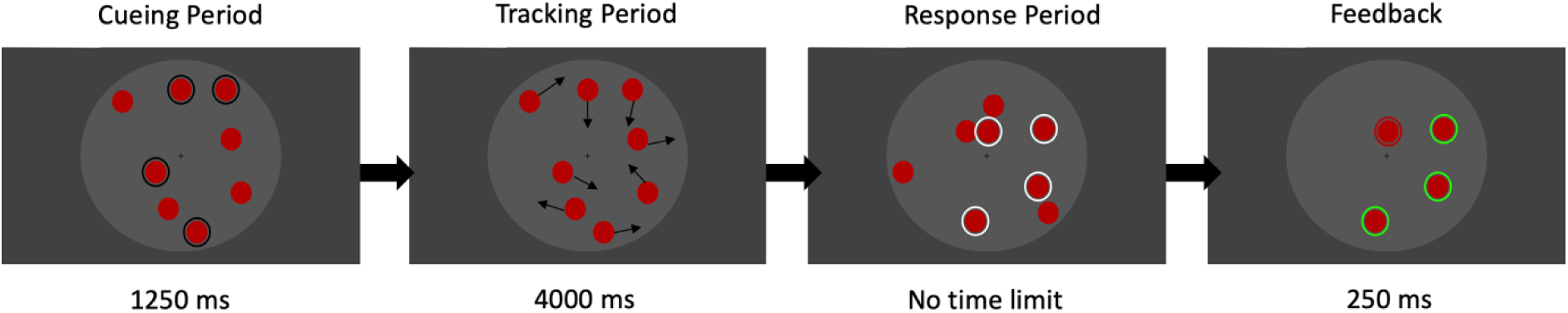
Illustration of the trial sequence in Experiment 1. Targets and distractors flickered at designated frequencies throughout cueing and tracking phases. Targets and distractors moved at one of three speeds during the tracking phase. Black arrows indicate motion directions and weren’t displayed. Participants responded using consecutive mouse clicks and could select or deselect objects as needed. They could only proceed to the next trial once exactly four objects were selected.

Throughout each trial, half of the discs flickered at 10 Hz whilst the other half flickered at 12 Hz. Depending on the condition, the 10 Hz discs served as targets and 12 Hz discs as distractors, or vice versa.

The experiment comprised 336 trials across eight blocks of 42 trials each resulting in 56 trials per condition. The three speed conditions were duplicated for the two target frequencies, yielding six conditions overall. Each block of 42 trials contained exactly 7 trials from each condition, which were presented in random order. A set of 56 starting positions and initial motion directions of objects were generated and copied across the six conditions (in randomized order) to minimize physical stimulus differences between conditions.

#### Behavioral data analysis

Responses in each trial were classified as correct if all four targets were reported correctly. Accuracy rates were analyzed by means of a one-way repeated measures ANOVA with factor Speed. All ANOVA analyses used Greenhouse-Geisser correction for non-sphericity where applicable.

#### EEG acquisition and preprocessing

EEG data were recorded using an ActiveTwo amplifier system (Biosemi) from 64 Ag/AgCl electrodes at 256 Hz sampling rate. The manufacturer’s standard electrode montage was modified by relocating electrodes from T7/8 and F5/6 positions to PO9/10 and I1/2 to enhance spatial coverage over occipital regions. Electrooculographic activity was monitored using electrodes positioned above and below the left eye (vertical EOG) and at the outer canthi of both eyes (horizontal EOG). Data processing was conducted using the EEGLAB toolbox (Delorme & Makeig, 2004) and custom MATLAB (MathWorks Inc., Natick, MA) routines.

Epochs spanning 400 to 3900 ms post-motion onset were extracted from the continuous data. Trials containing blinks or eye movements exceeding 20 μV were rejected. Each epoch was detrended by removing the epoch mean and linear trend. The pre-processed data were then submitted to an automated artefact detection routine (Junghöfer et al., 2000) that either replaces contaminated sensors using statistically weighted spherical interpolation or rejects entire trials when excessive sensors are compromised.

After removing data from participants with excessive EEG artifacts or poor behavioral performance, the average trial rejection rate was 20.1% (±9.4%), whilst the average number of interpolated channels was 3.83 (±1.24).

#### SSVEP Analysis

SSVEP analysis was restricted to trials with correct behavioral responses, yielding an average of 36 epochs per condition per participant (±7.05) out of the total of 56 trials per condition. Epoched data underwent scalp current density (SCD) transformation via spherical spline interpolation (Perrin et al., 1989). Following the preregistration protocol, SSVEP amplitudes at the target frequencies (10 Hz and 12 Hz) were extracted from SCD-transformed epochs as the absolute values of complex Fourier coefficients for each frequency, condition, and participant. Prior to statistical analysis, SSVEP amplitudes for each electrode were averaged for each condition, participant and frequency and rescaled by dividing each of the resulting amplitudes by the mean over all conditions (Adamian & Andersen, 2024). The resulting amplitudes scaled to the mean of 1.0 were combined across frequencies to produce average SSVEP amplitudes for the three speed (Experiment 1) or density (Experiment 2) conditions in each cluster of electrodes. These amplitudes were submitted to a repeated measures ANOVA with factors Attention (Attended vs Unattended) and Speed (3.8 dva/s, 7.6 dva/s and 10.9 dva/s).

#### Deviation from preregistration

Following the preregistration protocol, SSVEP amplitudes were calculated at midline occipital electrodes (Oz and POz). However, it was later decided to repeat the analysis using lateral occipital electrodes (P7/8 and PO7/8). This cluster of electrodes exhibited strong SSVEP signal, in line with a previous study using a similar paradigm (Adamian & Andersen, 2022). This study demonstrated that attentional enhancement of SSVEPs specifically in the lateral cluster of electrodes was associated with higher probability of successful tracking, suggesting particular relevance of these scalp locations for multifocal attention to moving objects.

### Results

Participants made more errors when tracking faster moving objects (*F*_(2,32))_ = 37.58, *p* < 10^−8^, *η^2^* = 0.35). Tracking accuracy (Figure 2A) was reduced for the 7.6 dva/s condition compared to 3.8 dva/s (*t*_(33)_ = 6.53, *p* <10^−6^) and was lowest when tracking objects moving at 10.9 dva/s (*t*_(33)_ = 6.62, *p* < 10^−6^).

**Figure 2.**
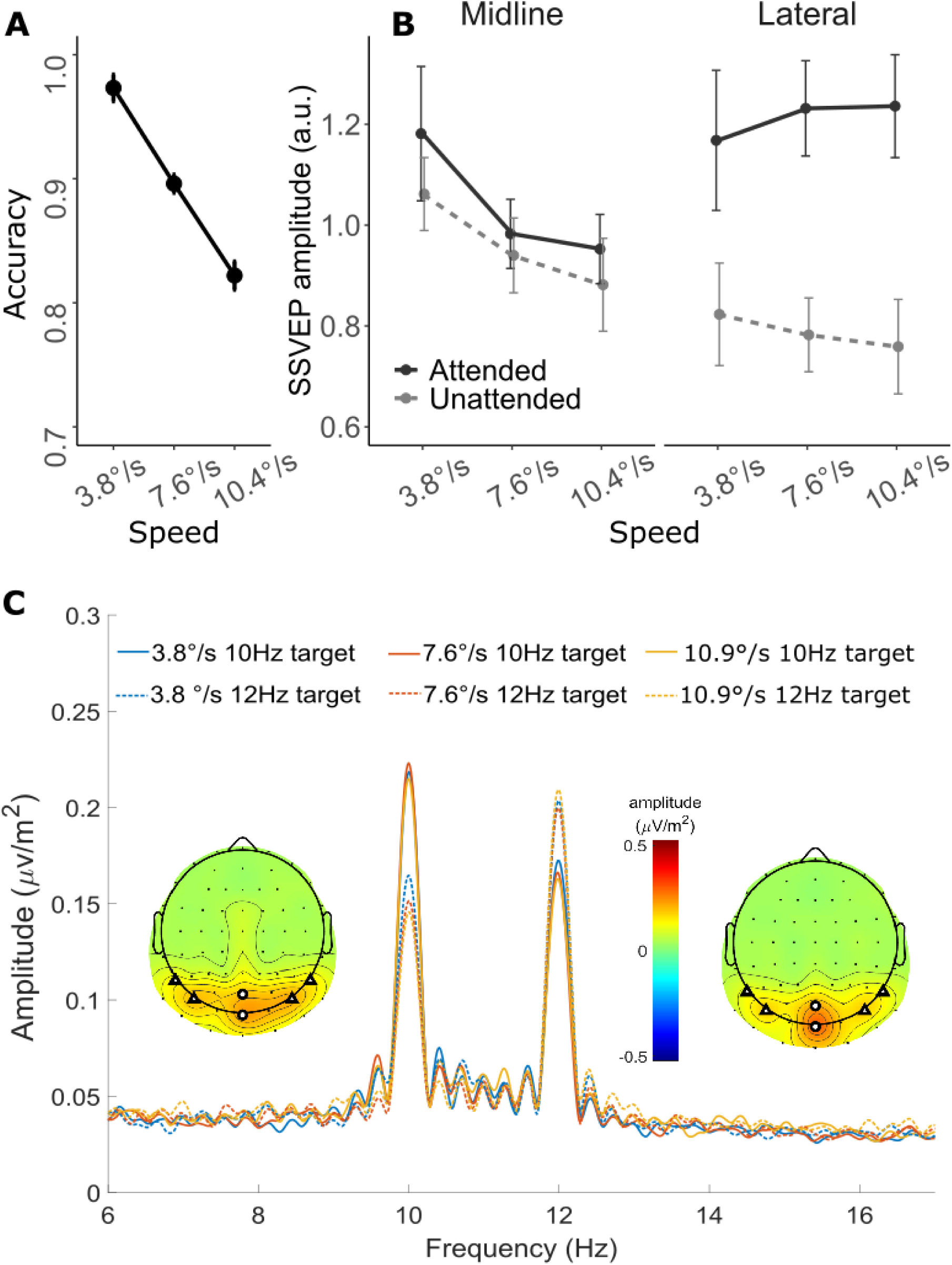
Results of Experiment 1. **A**: Mean accuracy rates for three speed conditions; **B:** Rescaled grand mean SSVEP amplitudes at midline and lateral electrode clusters. **C:** Grand-averaged amplitude spectrum of both electrode clusters combined, obtained by Fourier transformation and zero-padded to 16,384 points. Inset: Grand mean scalp current density (SCD) maps of SSVEP amplitudes at 10 Hz (left) and 12 Hz (right) averaged across conditions. Circles on topographical maps indicate Midline cluster electrodes, triangles indicate Lateral cluster electrodes. Error bars denote within-subject 95% confidence intervals (Morey, 2008)

#### SSVEP amplitudes

In the midline cluster of electrodes (Figure 2B left), SSVEP amplitudes elicited by attended objects were higher than those elicited by unattended objects (*F*_(1,16)_ = 5.08, *p* = 0.039, *η^2^* = 0.05, BF₁₀ = 1.25), confirming that attention facilitated processing of tracked targets in early visual cortex. SSVEP amplitudes decreased with increased object speed (*F*_(2,32)_ = 6.58, *p* = 0.004, *η^2^* = 0.22, BF₁₀ = 1872) and there was no interaction between speed and attention (*F*_(2,32)_ = 0.71, *p* = 0.497, *η^2^* = 0.07, BF₁₀ = 0.18).

Because attentional modulation of SSVEP amplitudes is multiplicative rather than additive (Adamian & Andersen, 2024), the presence of the main effect of speed precludes direct interpretation of the interaction term. To assess whether attention effects varied across speed conditions, we therefore computed an attentional modulation index (AMI) for each condition separately, and entered AMIs into a one-way ANOVA with factor Speed:

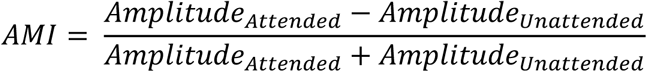

This additional analysis confirmed that attention was not modulated by speed (*F*_(2,32)_ = 0.43, *p* = 0.65, *η^2^* = 0.01, BF₁₀ = 0.21).

In the lateral cluster of electrodes (Figure 2B right), SSVEP amplitudes were not modulated by object speed (*F*_(2,32)_ = 0.06, *p* = 0.946, *η^2^*= 0.002, BF₁₀ = 0.09), but they were strongly enhanced by attention (*F*_(1,16)_ = 52.34, *p* < 10^−6^, *η^2^* = 0.38, BF₁₀ = 2.25 × 10¹⁶). The interaction between speed and attention was not significant (*F*_(2,32)_ = 1.55, *p* = 0.228, *η^2^*= 0.015, BF₁₀ = 0.28), indicating that the magnitude of attentional facilitation was stable across tracking speeds.

### Discussion

The results of Experiment 1 suggest that the strength of attentional selection of successfully tracked targets does not depend on their speed, as we observed no decrease in the relative attentional enhancement of targets compared to distractors with increased object speed in either electrode cluster. Importantly, behavioral performance was affected – participants tracked faster moving objects less successfully. The magnitude of this performance decrease is in line with the previous study where the number of tracked objects was manipulated (Adamian & Andersen, 2022). However, despite similar behavioral effects, only one of these two bottlenecks of tracking performance affects distribution of attention. While increasing the number of tracked targets weakens attentional facilitation each target receives, increasing their speed does not produce corresponding changes in attentional distribution as indexed by SSVEPs. Tracking errors due to speed therefore do not stem from weakened selection, but rather from bottom-up display characteristics that challenge this selection.

The flexible-resource model proposes that faster moving targets inherently require more attentional resources (Alvarez & Franconeri, 2007) leading to a smooth trade-off between the number of objects that can be tracked and their speed. However, it is unclear whether this trade-off results from less effective allocation of attention to faster targets or whether the same amount of attention becomes insufficient to track objects at higher speeds. In our paradigm, only speed was manipulated whilst spatial arrangement and trial duration remained constant, inevitably increasing the number of close encounters at higher speeds. Yet SSVEPs in successful tracking trials did not vary systematically with speed, suggesting that the strength of attentional selection remained constant despite increased bottom-up interference. This aligns with findings from Drew et al. (2013) who manipulated speed whilst examining tracking-related ERPs. Although their focus was on error trials, they found no effect of speed on the contralateral delay activity (CDA) when participants tracked a single target, further suggesting that speed does not modulate the attentional resources allocated to tracked objects.

Interestingly, we observed that attentional modulation was weaker in the midline electrode cluster compared to lateral clusters, and SSVEP amplitude decreased with speed specifically in the midline cluster. This pattern may reflect differences in receptive field sizes across the visual hierarchy (Smith et al., 2001). Midline electrodes likely reflect activity from earlier visual areas (V1-V3), which have smaller receptive fields than higher-order areas such as V4 or MT represented by lateral electrode clusters. At faster speeds, objects transit more rapidly through these smaller receptive fields, spending less time within any single receptive field and thereby producing weaker SSVEPs. In contrast, the larger receptive fields of higher visual areas may be better able to maintain continuous representation of fast-moving objects, resulting in speed-invariant attentional modulation in lateral clusters.

One alternative explanation for the speed invariance of attention in this study is that participants could, knowingly or unknowingly, select targets based on their common flicker frequency rather than the cue. To rule out this possibility, we carried out a separate behavioral control experiment with 17 different participants. This experiment used the same MOT paradigm as the main study but manipulated whether targets and distractors flickered at the same frequency or at two different frequencies. If participants were able to utilize flicker frequency as a cue for tracking, performance should be better when targets and distractors flicker at different frequencies. Participants tracked four targets amongst eight total objects moving at one of three speeds, identical to Experiment 1. In the critical manipulation, objects either 1) all flickered at 10 Hz, 2) all flickered at 12 Hz, 3) targets flickered at 10 Hz whilst distractors flickered at 12 Hz, or 4) vice versa. For analysis, these conditions were collapsed into two levels: frequency-same (conditions 1-2) and frequency-different (conditions 3-4). While tracking accuracy was lower at higher speeds (*F*_(2,32)_ = 29.99, *p* < 10^−7^, *η^2^* = 0.28), it did not depend on whether all targets were flickering at the same frequency or not (Frequency: *F*_(1,16)_ = 2.89, *p* = 0.109, *η^2^* = 0.04; Speed x Frequency interaction: *F*_(2,32)_ = 1.96, *p* = 0.157, *η^2^* = 0.007). This indicates that participants were unable to use differences in flicker frequency as a cue for attentional selection, consistent with previous studies (Müller et al., 2006; Störmer et al., 2013).

## Experiment 2: Total Number of Objects

Having established that attentional selection of tracked targets remains robust across different speeds, we next examined whether increasing the total number of objects in the display dilutes attention distributed among a fixed number of tracked targets. This study is similar to Experiment 1, but instead of tracking targets at different speeds, participants tracked four targets among 8, 12, or 16 objects. Other spatial parameters of stimulation such as object sizes or total area remained unchanged. According to the flexible resources account, increased number of close encounters between targets and distractors depletes attention through higher demands on spatial resolution (Alvarez & Franconeri, 2007; Iordanescu et al., 2009). If this was the case, we expect to see weakened attentional selection with increased number of distractors.

### Method

#### Participants

Forty-eight University of Aberdeen students were recruited to participate in the study (10 men, 37 women, 1 non-binary; 2 left-handed; M_age_ = 21.38, SD_age_ = 6.46). Five of them participated in the previous study. There were two reasons for the increased sample size compared to Experiment 1. First, we wanted to increase statistical power to detect any possible small interactions with attention. Second, we planned an analysis of individual differences whose purpose is orthogonal to the present paper, and which will be published separately. All participants reported normal or corrected-to-normal visual acuity and normal color vision. The study received ethical approval from the Psychology Ethics Committee at the University of Aberdeen (No. 4412909).

Data from seven participants were excluded using the same exclusion rules as Experiment 1. The final sample consisted of 41 participants.

#### Stimuli and procedure

Experiment 2 employed the same MOT paradigm but manipulated the total number of stimuli rather than their speed (Figure 3). Participants tracked four targets amongst either 8, 12, or 16 total objects, all moving at a constant speed of 5.7 dva/s. The speed was determined during the piloting stage to avoid ceiling effects in behavioral performance. To alleviate spatial constraints within the fixed viewing area the discs and the invisible surrounds was slightly reduced in size (3.8 dva diameter disks with additional 0.8 dva “bumpers”). The full report procedure yielded equal guessing chance across speed conditions in Experiment 1, however this would not have been the case with respect to object number conditions in Experiment 2 as the proportion of targets in the display is systematically confounded with the total number of objects. Therefore, the response procedure was changed as follows. After each trial, two discs were consecutively probed, with targets and distractors probed with 50% probability. Participants were asked to report by key press whether each of the probed disks was a target or a distractor. This ensured comparable chance performance levels across conditions and experiments, thereby facilitating comparisons.

**Figure 3.**
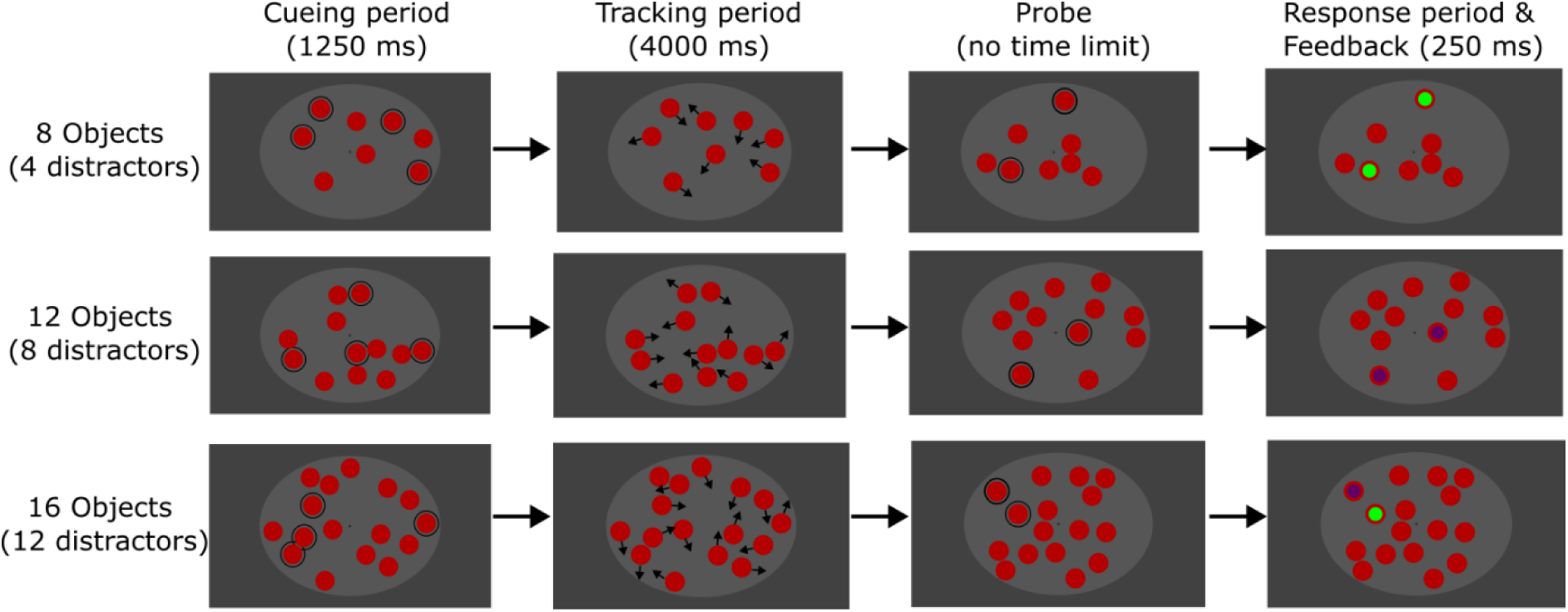
Illustration of the trial sequence in Experiment 2. Probes were presented and responses collected consecutively. Feedback displayed for both correct responses (top), both incorrect responses (middle) and one correct response (bottom). Black arrows during tracking period indicate motion directions and weren’t displayed.

Irrespective of condition, throughout the trial four discs flickered at 10.9 Hz and four discs flickered at 13.3 Hz. In conditions where more than eight discs were presented, the remaining discs flickered at 12 Hz. To-be-tracked discs always included all four 10.9 Hz or all four 13.3 Hz discs allowing us to compare attentional enhancement of targets relative to distractors regardless of the total number of objects in the set. There were 336 trials in total with 56 trials per condition (three object number conditions for each of the two target stimulation frequencies).

After the EEG session, participants also completed the Adult Self-Report ADHD Scale (Kessler et al., 2005) and the Mind Excessively Wandering Scale (Mowlem et al., 2019), the results from which are a part of a separate project not reported in this manuscript.

#### Data analysis

Behavioral and EEG data analysis followed the same processing and analysis pipeline as Experiment 1. Only trials with correct behavioral responses (to both probed stimuli) were included in the analysis. After removal of EEG artifacts and epochs preceding incorrect responses, the average trial rejection rate was 27.8% (±16.0%) with the 3.83 (±1.24) channels interpolated. Rescaled SSVEP amplitudes were analyzed by means of repeated measures ANOVA with factors Attention (attended vs unattended) and Number of objects (8, 12, and 16).

### Results

Tracking accuracy decreased in line with the number of stimuli on the screen (*F*_(2,80))_ = 63.23, *p* < 10^−16^, *η^2^* = 0.06). Pairwise comparisons confirmed that tracking accuracy decreased when the total number of objects increased from 8 to 12 (*t*_(40)_ = 7.59, *p* <10^−8^, *d* = 0.84) and further decreased from 12 to 16 objects (*t*_(40)_ = 5.48, *p* <10^−5^, *d* = 0.60).

#### SSVEP amplitudes

In the midline cluster of electrodes (Figure 4B) SSVEP amplitudes were significantly higher for attended compared to unattended objects (*F*_(1,40)_ = 21.55, *p* < 10^−4^, *η^2^* = 0.18 BF₁₀ = 6.91 × 10⁸). SSVEP amplitudes were significantly modulated by the total number of objects (*F*_(2,80)_ = 5.24, *p* = 0.007, *η^2^* = 0.046, BF₁₀ = 3.3). Crucially, there was no significant interaction between the number of objects and attention (*F*_(2,80)_ = 0.97, *p* = 0.38, *η^2^* = 0.005, BF₁₀ = 0.11), suggesting that the magnitude of attentional facilitation did not depend on the total number of objects in the set.

**Figure 4.**
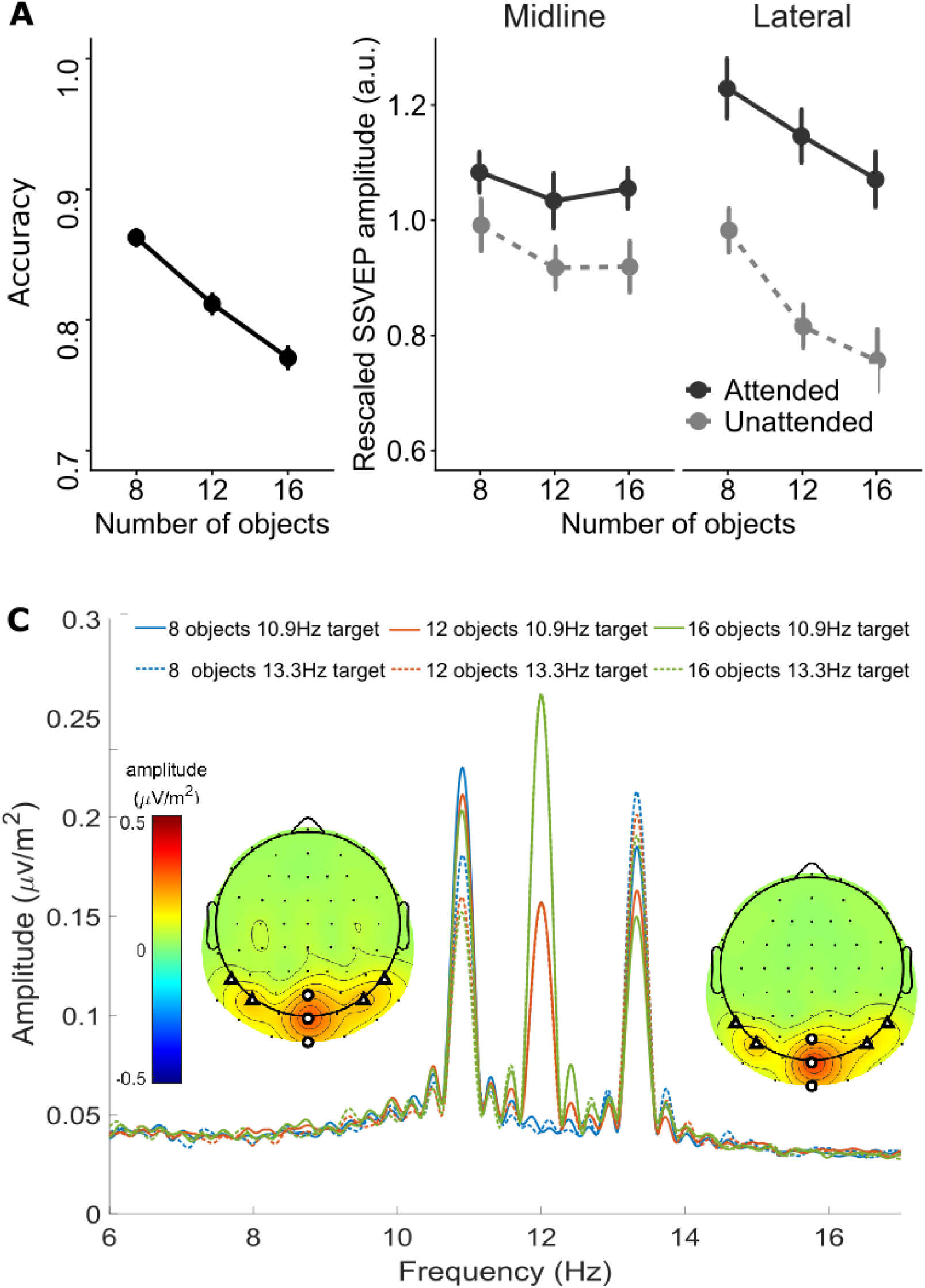
Results of Experiment 1. **A**: Mean accuracy rates for three density conditions; **B:** Rescaled grand mean SSVEP amplitudes and midline and lateral electrode clusters. **C:** Grand-averaged amplitude spectrum of both electrode clusters combined, obtained by Fourier transformation and zero-padded to 16,384 points. The 12 Hz peak does not appear in 8 object conditions as no objects flickered at 10 Hz. Inset: Grand mean scalp current density (SCD) maps of SSVEP amplitudes at 10.9 Hz (left) and 13.3 Hz (right) averaged across conditions. Circles on topographical maps indicate Midline cluster electrodes, triangles indicate Lateral cluster electrodes. Error bars denote within-subject 95% confidence intervals (Morey, 2008)

In the lateral cluster of electrodes, attention also enhanced SSVEP amplitudes (*F*_(1,40)_ = 107.32, *p*<10^−12^, *η^2^* = 0.54, BF₁₀ = 1.32 × 10^31^). Amplitudes in both attended and unattended conditions decreased with the total number of objects (*F*_(2,80)_ = 42.82, *p* < 10^−9^, *η^2^* = 0.25, BF₁₀ = 3607341). There was no interaction between the two factors (*F*_(2,80)_ = 2.63, *p* = 0.08, *η^2^* = 0.017, BF₁₀ = 0.40).

Following the same analysis logic as the first experiment, to assess whether attention effects varied across conditions in the presence of the main effect of object number, we computed AMI for each condition separately. Contrary to the prediction of attention weakening, in the lateral cluster AMIs increased slightly with increasing number of objects (8 objects: M = 0.11, SD = 0.09; 12 objects: M = 0.17, SD = 0.08; 16 objects: M = 0.18, SD = 0.09; *F*_(2,80)_ = 5.81, *p* = 0.004, *η^2^* = 0.05). AMIs in the midline cluster were not modulated by the number of objects (*F*_(2,80)_ = 1.09, *p* = 0.342, *η^2^* = 0.009).

### Discussion

Increasing the number of objects in the display did reduce tracking performance as expected (Bettencourt & Somers, 2009; Drew et al., 2013; Feria, 2012), although this effect was less pronounced than the performance decrements observed with speed manipulations in Experiment 1 (see Figs. 2a & 3a), and in our previous work manipulating the number of targets to be tracked (Adamian & Andersen, 2022). This was not accompanied by a corresponding drop in attentional modulation of SSVEPs, which even exhibited a slight increase with increasing object numbers in the lateral electrode cluster. Therefore, increased tracking difficulty with more objects was not mediated by less effective attentional allocation.

Our results can be interpreted from the perspective of the biased competition model of attention (Desimone & Duncan, 1995) according to which simultaneously presented stimuli activate overlapping neural populations and compete for cortical representation. This competition is affected by bottom-up (stimulus) and top-down (attentional) factors, biasing neural responses in favor of attended stimuli therefore allocating limited cortical resources to task-relevant objects (Luck et al., 1997; Reynolds & Chelazzi, 2004; Reynolds & Heeger, 2009). The competition for neuronal representation is strongest when stimuli fall into shared receptive fields and attention affects neuronal firing rates most when this is the case (Moran & Desimone, 1985). This explains why the lateral cluster exhibits a more pronounced drop in SSVEP amplitudes with increasing object numbers and stronger attentional modulation than the midline cluster, as it reflects later visual areas with larger receptive fields and therefore stronger competition. As more objects are added to the display, they act in competitive suppression, reducing neural response to any individual object (Kastner et al., 1998, 2001). Note that according to biased competition, the intensified competition for neural representation from adding more objects may lead to larger attentional modulation of visual processing, even if the attentional top-down signals themselves remain unchanged (however, Andersen, Müller, & Martinovic (2012) and Keitel et al. (2013) reported independent effects of competition and attention on SSVEP amplitudes).

Attention might facilitate tracking by reducing crowding between targets and distractors. Cueing spatial attention can decrease crowding effects (Yeshurun & Rashal, 2010). Some authors have suggested that crowding arises due to a limitation of precision of attention allocation (Intriligator & Cavanagh, 2001). If crowding was a limiting factor and this interpretation of crowding was true, then we should observe reduced attentional allocation in the condition with more distractors. However, this is not what we observed.

Consistent with Experiment 1, the dissociation between declining behavioral performance and maintained or enhanced attentional modulation demonstrates that tracking failures with increased object speed and number are not mediated by less effective attentional allocation. Similarly to limitations on tracking faster objects, spatial interference increases the probability of tracking errors even though the strength of attentional modulation is maintained.

#### Template-based source localization of attention effects

The analyses carried out for Experiments 1 and 2 relied on electrode selection based on SSVEP amplitudes at the scalp level. Previous studies from our lab applying surface current density (SCD) transformation to SSVEP data have repeatedly identified independent electrode clusters whose activity was differentially modulated by selective attention (Adamian et al., 2020; Adamian & Andersen, 2022; Andersen et al., 2012). In the current study, our experimental manipulations also affected the two clusters in distinct ways: midline cluster amplitudes were decreased with increasing object speed, whilst lateral cluster amplitudes were decreased with increasing object density. However, this method of electrode selection does not reveal the specific neural sources underlying these scalp-recorded signals. While we have anatomic grounds to assume that midline electrodes primarily reflect V1-V3 activity and lateral electrodes reflect activity in MT/V5/LOC, source localization can provide a more granular view of the cortical origins of attention effects.

To source localize attention effects, we applied a novel approach that uses fMRI-defined templates which specify how activity of different visual cortical areas appear on the scalp and therefore, in EEG data (Poncet & Ales, 2023). Templates created by Poncet and Ales (2023) are based on MRI and fMRI data of 50 participants and were generated by identifying retinotopic visual areas with fMRI, then simulating the expected EEG scalp distribution for each area. Templates are then fit to the observed EEG data using regularized regression to determine the contribution of each brain source to the recorded scalp activity. The templates and the accompanying EEG Source Template Matching toolbox are openly available.

#### Method

Source localization was performed using EEG templates corresponding to average scalp activity for 18 visual cortical areas created by Poncet and Ales (2023). EEG data were referenced to the average. Complex Fourier coefficients were averaged across participants and submitted to the template matching algorithm, resulting in estimated activity for each condition and each region of interest (ROI). Since the template matching method uses group-level data, statistical tests were based on bootstrap resampling with replacement (1000 iterations).

For each bootstrap sample, AMIs were calculated separately for each ROI, then averaged across bilateral ROIs and across the two stimulation frequencies to reduce multiple comparisons. Additionally, AMIs were averaged between ventral and dorsal V2 and V3 to account for full-field stimulation, resulting in 7 ROIs in total: V1, V2, V3, V4, V3A, LOC and hMT+. To identify cortical ROIs significantly modulated by attention, we performed two-sided bootstrap t-tests of each condition-average AMI against zero.

#### Results

In Experiment 1 where object speed was manipulated, six out of seven visual ROIs showed significant modulation of SSVEP amplitudes by attention (see Figure 5; *p*s < 0.001), with the exception of area V1 *(p* = 0.26). In Experiment 2, where total number of objects varied by condition, significant attentional modulation was observed in V2 (*p* = 0.02), V4

**Figure 5.**
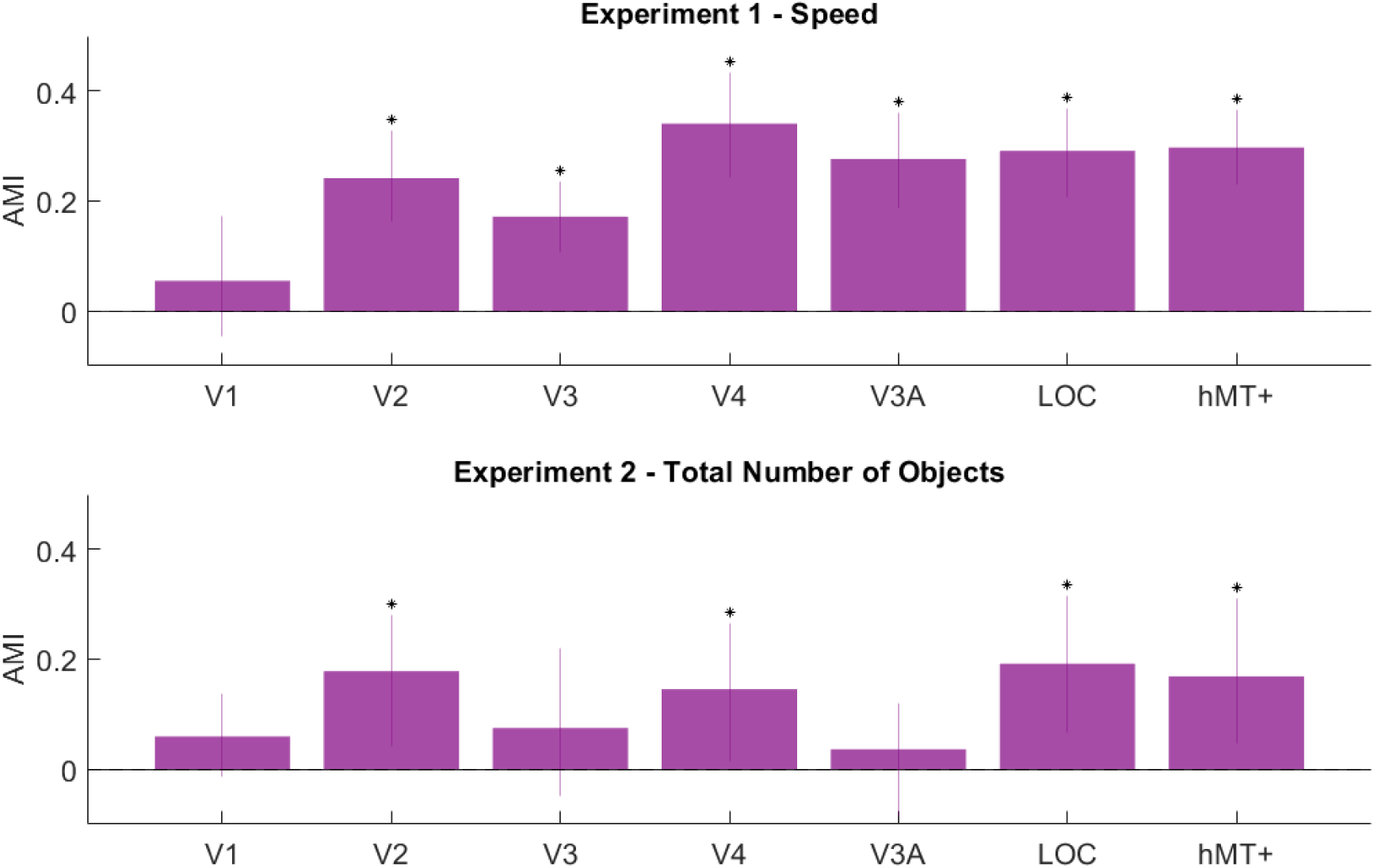
Results of the source localisation analysis for Experiment 1 (top) and Experiment 2 (bottom). Error bars are 95% Cis. Stars indicate statistically significant AMIs.

(*p* = 0.03), LOC (*p* = 0.01) and hMT+ (*p* = 0.01). Again, SSVEP signal originating in V1 was not modulated by attention (*p* = 0.08). The absence of significant attentional modulation in V1 may reflect its small receptive fields, which cannot integrate SSVEPs as tagged objects move through space. This likely results in weak or unstable SSVEP responses in V1, leaving little signal to be enhanced by attention. While V1 along with MT is an established major source of SSVEP (Di Russo et al., 2007), it is unsurprising that the use of moving stimuli would shift the balance of SSVEP amplification towards extrastriate visual areas.

Areas V3 and V3A showed attentional modulation only in the speed manipulation experiment. While speculative, this could reflect enhanced motion processing associated with successful tracking of faster moving objects (Tootell et al., 1997).

These results are in broad agreement with previous studies identifying peak modulation of SSVEP amplitudes in early visual cortex (V1-V3) and MT/LOC (Adamian et al., 2020; Störmer et al., 2013), validating template matching as an effective method for localizing attention effects in SSVEP experiments. Compared to phase-based electrode clustering, this approach provides finer anatomical resolution by separating signals from adjacent visual areas that could otherwise be conflated in electrode-level analyses.

## General Discussion

In two experiments, we investigated if object speed and number of distractors modulate attentional allocation during multiple object tracking in the same manner as top-down task demands. In Experiment 1, we manipulated object speed whilst holding the number of targets and distractors constant. Behaviorally, tracking performance declined as speed increased, replicating well-established findings (Drew et al., 2013; Feria, 2013). However, SSVEPs revealed that attentional modulation of tracked targets remained constant across speeds during successful tracking trials. A behavioral control experiment further confirmed that the different flicker frequencies between targets and distractors did not benefit tracking performance. In Experiment 2, we manipulated the total number of objects in the display whilst keeping the number of targets constant. Again, tracking performance declined as more objects were added, yet attentional modulation either remained constant (midline electrode cluster) or slightly strengthened (lateral electrode cluster) with increased object number. These findings contrast sharply with the effects of tracking more targets: when the number of tracked targets increases, attentional modulation of each individual target systematically weakens (Adamian & Andersen, 2022). Together, these results demonstrate that bottom-up display characteristics and top-down task demands affect tracking in fundamentally different ways. Although these manipulations may trade-off behaviorally (Alvarez & Franconeri, 2007), they have qualitatively different effects on visual cortical processing of targets and distractors. Consequently, even if conditions in which different numbers of targets tracked are matched in tracking performance by covarying physical stimulus parameters, this may result in perceptual or behavioral differences in other measures than tracking performance.

This dynamic can be explained using a classic metaphor of sticky references (Pylyshyn, 1989), where attention is thought of as “glue” binding targets to their representations in visual cortex. When tracking more targets, the same amount of glue must be spread more thinly, and each target receives less attentional facilitation (Adamian & Andersen, 2022). However, with increased bottom-up demands such as speed or number of targets, the amount of glue stays the same while external forces pulling targets away from their representations (e.g. close encounters) become stronger and/or more frequent. The amount of glue (attentional allocation) remains constant, but it becomes insufficient to maintain stable tracking against heightened interference, resulting in more errors. In this view, there is no gradual weakening of attention in physical stimulus conditions that are more challenging for tracking. However, attentional selection may break down for individual objects when discrete tracking errors occur, such as the loss of a target object (“dropping”) or a close encounter between objects leading to erroneous tracking of a distractor object instead (“swapping”) (Drew et al., 2013). For this reason, SSVEP amplitudes during tracking predict behavioral performance at the end of the trial (Adamian & Andersen, 2022; Störmer et al., 2013).

The pattern of results supports target enhancement rather than distractor suppression as the primary attentional mechanism subserving object tracking (Bettencourt & Somers, 2009). If tracking relied on active suppression of distractors, one would predict that attentional modulation should decrease with increasing numbers of distractors. However, both the current findings and our previous work (Adamian & Andersen, 2022) demonstrate the opposite pattern: attentional modulation is sensitive to the number of tracked targets but remains stable across variations in distractor number. This dissociation strongly suggests that the visual system prioritises enhancing target representations rather than suppressing distractor processing during multiple object tracking.

The dissociation between top-down attentional control and bottom-up task demands is consistent with prior evidence. Neuroimaging studies have revealed functionally distinct neural networks for general task engagement versus tracking specific numbers of targets (Culham et al., 2001; Felßberg et al., 2025; Mäki-Marttunen et al., 2020). Behavioural work has similarly suggested that spatial interference imposes limits on tracking that are separable from attentional capacity limits (Meyerhoff et al., 2016; Vul et al., 2009). The present findings add to this evidence by demonstrating that manipulations affecting bottom-up processing demands such as object speed or distractor number are orthogonal to top-down attentional selection during successful tracking.

To conclude, while tracking is accomplished through multifocal attention, it is not less effective allocation of selective attention that limits tracking capacity under high-speed or high-distractor-number conditions. Attention is allocated just as effectively but does not afford the same level of tracking performance under these more challenging conditions.

## Acknowledgements

We thank Rosalind Hillhouse and Thomas Alexander for assistance with data collection, and Marlene Poncet and Justin Ales for guidance in applying template matching analysis to SSVEP data. This work was supported by Leverhulme Early Career Fellowship ECF-2020-488 awarded to NA.

## Declarations

### Ethics approval

This study was performed in line with the principles of the Declaration of Helsinki. Approval was granted by the Psychology Ethics Committee at the University of Aberdeen (No. 4412909 and PEC/5048/2022/7.

### Consent to participate

Informed consent was obtained from all individual participants included in the study.

### Consent for publication

As part of informed consent procedure, participants were informed and agreed for their anonymised data to be shared via an online repository.

### Availability of data and materials

Participant-level data (EEG and behavioural) and aggregated data with analysis code is available: https://osf.io/ypgfs

### Code availability

Code producing data visualization and statistical analyses is available (https://osf.io/ypgfs).

### Authors’ contributions

**Conceptualization:** N.A. and S.K.A.

**Data curation:** N.A. and F.A.

**Funding acquisition:** N.A. and S.K.A.

**Investigation:** N.A., F.A., and S.K.A.

**Methodology:** N.A., F.A., and S.K.A.

**Project administration:** N.A. and F.A.

**Resources:** N.A.

**Software:** N.A.

**Supervision:** N.A. and S.K.A.

**Visualization:** N.A.

**Writing - original draft:** N.A. and F.A.

**Writing - review & editing:** N.A. and S.K.A.

